# Resource co-limited growth in fluctuating environments

**DOI:** 10.1101/356980

**Authors:** Apostolos-Manuel Koussoroplis, Svenja Schälicke, Michael Raatz, Moritz Bach, Alexander Wacker

**Affiliations:** Theoretical Aquatic Ecology and Ecophysiology group, Institute of Biochemistry and Biology, University of Potsdam, Potsdam, Germany.; L.M.G.E. UMR CNRS 6023, Université Clermont Auvergne, Aubière, France.

**Keywords:** cholesterol, co-limitation, covariance, Daphnia, digestive acclimation, dynamic energy budgets, environmental autocorrelation, food quality, phosphorus, storage, stressor coincidence, temporal ecology, unbalanced diets

## Abstract

Variability in the supply of limiting resources determines consumer-resource interactions. Yet, how consumers are affected by variability when multiple resources co-limit growth remains unknown. We use a two-resource DEB model to predict how consumer somatic growth rate responds to the temporal structure (i.e. fluctuation frequency, phase and covariance) of single and co-limiting resources supply. Subsequently, we experimentally test the model predictions using *Daphnia magna* (co-)limited by dietary phosphorus and cholesterol supply. Both model and experiments indicate that for certain fluctuation frequencies, resource fluctuation phase and (co)variance can heavily affect somatic growth. The model suggests that dynamic resource storage and assimilation efficiency adjustment are key for predicting the frequencies at which the growth rate is mostly affected by (co)variance and phase. In a context of ongoing anthropogenic landscape homogenization, our results offer novel insights on how co-occurring perturbations to the temporal structure of resource supply can affect consumer performance.

## Introduction

Climate change is expected to increase the frequency of extreme events (Meehl 2004; Rahmstorf & Coumou 2011; Field & Intergovernmental Panel on Climate Change 2012). Providing local perturbations to established systems, these extreme events can promote environmental heterogeneity (Turner 2010). In contrast, anthropogenic impacts, such as pollution and industrialized agricultural practices, may induce biodiversity loss and simplify ecosystems thus decreasing environmental heterogeneity (Gámez-Virués *et al*. 2015). Within the tension of these two opposing scenarios the spatiotemporal variability perceived by organisms may either increase, decrease or remain unchanged.

The nutritional traits of plant or animal prey such as their composition of limiting resources or secondary metabolites strongly influence the organismal performance and consumer population dynamics (Sterner & Elser 2002; Simpson & Raubenheimer 2012; Hunter 2016; Sperfeld *et al*. 2016; Raatz *et al*. 2017). Spatiotemporal variability in the nutritional quality of food resources is inherent to natural consumer-resource systems (Orians & Jones 2001; Park *et al*. 2004; Junker & Cross 2014; Grosbois *et al*. 2016), and most consumers face intense and frequent fluctuations in food quality during their lifetime. Accumulating evidence shows that such resource variability strongly influences consumers at the individual (Stockhoff 1993; Hood & Sterner 2010; Pearse *et al*. 2018), population, and community level (Underwood 2004; Riolo *et al*. 2015) and may have far-reaching ecological implications, e.g. for the control of herbivore pest populations in agroecosystems (McArt & Thaler 2013; Wetzel *et al*. 2016). The importance of resource variability results from the deviation between performance achieved at fluctuating resource supply and performance achieved at constant, average resource supply (*variance effect*, Fig. 1A). If performance follows a non-linear saturating function of resource concentration, performance decreases for higher resource variability (Jensen’s inequality (Ruel & Ayres 1999; Wetzel *et al*. 2016)).

However, developing a more mechanistic basis of resource variability effects would allow better predictions of consumer population dynamics in natural communities.

**Figure 1:**
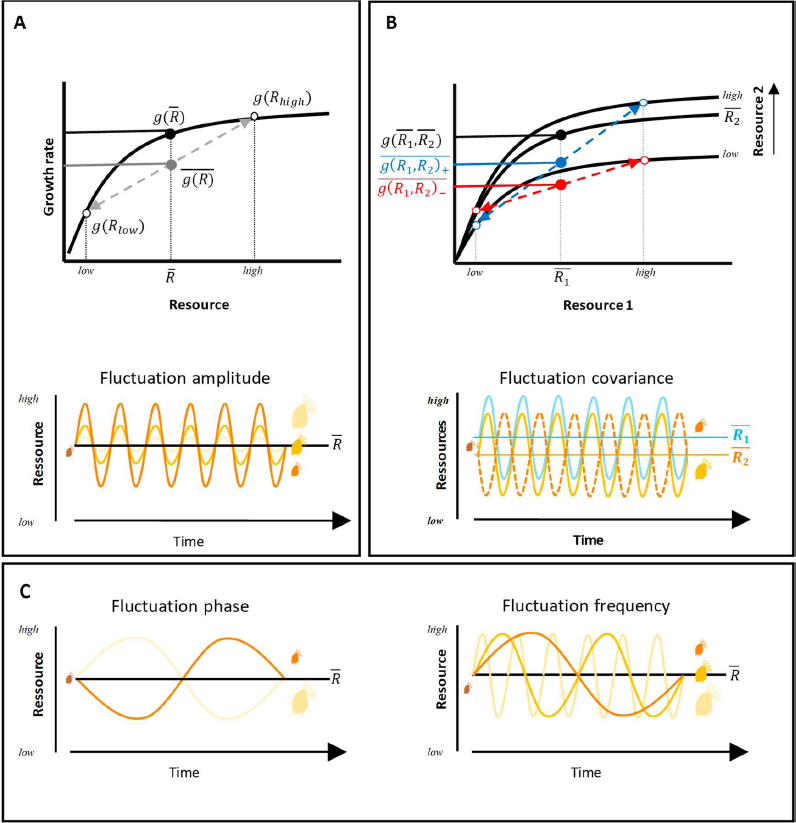
**Hypotheses** (**A**) For single resource limitation, non-linear averaging predicts that the average growth rate of a consumer experiencing fluctuating supply in a limiting resource R, 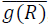, should be lower than the growth rate of a consumer experiencing a constant resource environment with same average resource conditions, 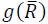. Consumer growth rate should decrease with increasing resource fluctuation amplitude 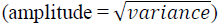 (**B**) When expanding to two synergistically co-limiting resources 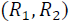, the average growth rate of a consumer experiencing positively covarying co-limiting resources, 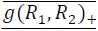, should be higher than that of a consumer experiencing negative covariance, 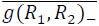. Note that an opposite covariance effect is expected if the co-limiting resources are antagonistic. (**C**) Because of reserve and acclimation effects (see text), the realized growth and therefore the exactitude of non-linear averaging predictions (see **A&B**) should be modulated by the resource fluctuation phase and frequency (for simplicity, only single resource cases are depicted).

To date, studies on resource variability focused on single nutritional traits (Carlotti *et al*. 2010; Hood & Sterner 2010; Wagner *et al*. 2017). Yet, consumer performance can be simultaneously driven by several nutritional traits of their food (e.g. concentrations of various mineral and biochemical resources or secondary metabolites), a situation known as co-limitation (Sperfeld *et al*. 2012, 2016). Hence, when feeding in nutritionally variable environments, consumers need to acquire, store and use multiple resources which might, or might not, co-occur temporally and spatially. For example, many consumers need to mix different food items with imbalanced nutrient ratios to acquire a sufficient blend of the co-limiting resources (Simpson & Raubenheimer 2012). Such consumers typically experience *negative covariance* in the co-limiting resources if the availability of the different food items does not coincide in time. Co-limiting resources can also co-occur at high and balanced concentrations in only one food item while being low or absent in all other. Ecological factors such as patchiness, fear, territoriality, or physical constraints (Winder *et al*. 2004; Hebblewhite & Merrill 2009; Lasley-Rasher *et al*. 2011; Camp *et al*. 2015) may only allow intermittent access to the high quality food items, the rest of time being spent on low quality food. This leads to *positive covariance* in co-limiting resources since their availability for the consumer peaks and drops simultaneously over time.

When co-limiting resources act non-additively, the resource covariance pattern considerably influences consumer performance (Koussoroplis *et al*. 2017a). A consumer should therefore perform differently in positive and negative covariance scenarios although temporal means and variances of the co-limiting resources are the same (*covariance effect sensu* (Koussoroplis & Wacker 2016; Fig. 1B). Given the propensity of co-limiting resources to interact (Simpson & Raubenheimer 2012; Sperfeld *et al*. 2016), the importance of resource variability for consumers cannot be addressed without considering the interplay of co-limitation and the spatiotemporal covariance of the co-limiting resources in the landscape.

Temporal and spatial resource variability are coupled and both translate to a perceived temporal fluctuation. Independently of whether the consumer is sessile in a dynamic environment (e.g. bivalves in tidal estuarine systems) or mobile in a heterogeneous landscape (e.g. daily vertically migrating zooplankton), nutritional variance and covariance are always experienced by the consumer as temporal fluctuations in resource supply. Changes in spatial or temporal contrasts of resource availability influence the amplitude of perceived resource fluctuation, whereas the grain-size of the environment (i.e. size or duration of food patches), the motility and life span, as well as the foraging behavior of the consumer influence the frequency and phase of the perceived fluctuations.

The performance of a consumer in a nutritionally variable environment can be empirically predicted from the resource-dependent responses generated under constant conditions (Sperfeld *et al*. 2016) using non-linear averaging methods (Wetzel *et al*. 2016; Denny 2017; Koussoroplis *et al*. 2017a). Yet, non-linear averaging needs to be regarded as a null-hypothesis based on the assumption that performance measured over longer time scales (typically several days) is a good approximation of the instantaneous performance in a fluctuating environment (Dowd *et al*. 2015). Obviously, this is not always the case as various physiological processes, which are not apparent under constant conditions, can manifest in a fluctuating environment. In this case, consumer performance in a variable environment deviates from that predicted by non-linear averaging (Niehaus *et al*. 2012; Kingsolver & Woods 2016; Koussoroplis *et al*. 2017a). For example, most living organisms first store recently acquired energy and nutrients into a metabolically inactive pool (hereafter *reserves* (Kooijman 2010)), which are subsequently mobilized and used for maintenance, growth, and reproduction. Such reserves may buffer the effects of environmental resource fluctuations on performance (hereafter *reserve effect*) (Muller & Nisbet 2000; Fujiwara *et al*. 2003; Hood & Sterner 2010). Furthermore, many organisms dynamically acclimate to fluctuating resources by e.g. adjusting nutrient extraction and transport through gut enzyme modifications (Karasov *et al*. 2011; Koussoroplis *et al*. 2017b), thereby improving assimilation efficiency of the resource that becomes most limiting (Clissold *et al*. 2010; Urabe *et al*. 2018). Acclimation processes may also have negative consequences if such processes are costly (Wetzel & Thaler 2016) or when acclimation lags behind the changes in environmental conditions (Kingsolver & Woods 2016). If acclimation is too slow compared to the fluctuations of the environment, organisms will underperform most of the time relative to non-linear average predictions (Niehaus *et al*. 2012; Koussoroplis *et al*. 2017a). This *acclimation effect* thus improves or decreases performance in fluctuating environments. Because of their innate physiological constraints, reserves and acclimation may operate at certain temporal scales, yielding a scale dependence of consumer performance, i.e. a different response to high and low frequencies at the same fluctuation amplitude (Koussoroplis *et al*. 2017a). Reserves and acclimation create a dependence of instantaneous growth on the nutritional history of an organism not only in terms of frequency, but also in terms of the phase of resource fluctuations.

Here, we test the hypothesis that for the same average and a fixed amplitude of resource quality fluctuations, environments with different temporal structures (i.e. resource supply fluctuation, covariance, frequency and phase, Fig. 1C) lead to altered consumer performances that do not necessarily comply with non-linear averaging predictions. We use a two-resource Dynamic Energy Budget (DEB) model (Fig. 2) to predict growth of a consumer in an environment with constant or fluctuating resource supply and compared the predictions to explore how the temporal structure of the environment mediates growth differences under fluctuating resources. Model predictions were experimentally reproduced by rearing juvenile *Daphnia* magna at variable phosphorus (P) and cholesterol supply. Our study offers both theoretical and empirical evidence that the temporal structure of resource availability can strongly decrease consumer performance in a fluctuating environment and provides a framework for modeling such negative effects. Using our model to predict the outcomes of combined changes in resource supply fluctuation amplitude and frequency enables novel insights on how landscape homogenization might affect local consumer species.

**Figure 2:**
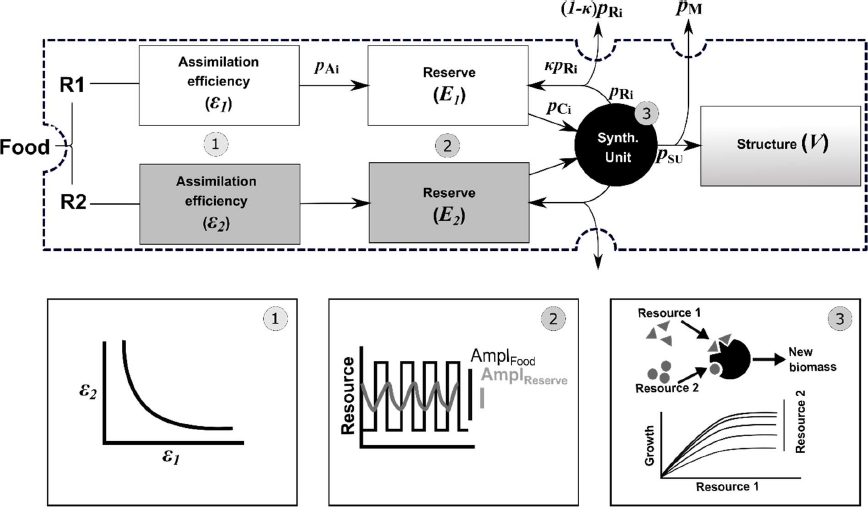
DEB model. Arrows indicate fluxes and rectangles indicate the state variables. Numbered circles indicate implemented physiological mechanisms: (i) The organism has the possibility to acclimate its assimilation efficiency to the resource that has the lower reserves (most limiting). The acclimation process involves a time lag and implies a trade-off shown in the left panel below. (ii) The organism has a reserve compartment for each resource that buffers the amplitude of higher frequency fluctuations in ingested resources (middle panel below). (iii) The synthesizing unit (SU) merges the resource fluxes that are liberated from the reserves into new biomass with a fixed resource ratio. The rate at which the SU produces biomass from the reserves depends on the intensity and the balance of the mobilized resources fluxes. The SU was implemented such that under constant resource conditions, the two resources act as interactively essential resources thereby enabling resource covariance effects.

## Materials and Methods

### Design of the study

Within this study the organism is exposed either to a sequence of resource regimes with high and low concentrations of two co-limiting resources, which fluctuate in different frequencies, or to different constant conditions. Five constant treatments result from supplying constant concentrations of both resources either high or low, one high and the other low or both at the mean of high and low (Fig. 1A-B; open and black circles). Constant treatments were used to calculate non-linear averaging predictions for growth of the consumer (Fig. 1A-B). In treatments of fluctuating resources, concentrations (high or low) are arranged in variance scenarios, where only one of the two resources fluctuates, and in positive and negative covariance scenarios with high concentrations of the two resources coinciding or alternating, respectively, keeping the same average resource concentration as in constant conditions treatments. Different fluctuation frequencies (i.e. time spans of the resource regimes) and different phases (i.e. initial combinations of resource concentrations) were studied. The total duration of exposure was the same across treatments. This allows comparing growth responses to constant and to fluctuating resource supply at different fluctuation frequencies (Fig. 1).

### Model structure

Based on General DEB Theory (Kooijman 2010) we constructed an ODE-model that predicts the growth of an organism (Tab. 1). The three important aspects of the model are (i) co-limitation by two interactively essential resources (this constitutes the null-model), (ii) independent storage of these resources in two reserves and (iii) regulation of assimilation efforts depending on the relative filling of the reserves with a trade-off between the two resources. The state variables of our model are the structural volume of the organism *V*, the respective resource density in the reserve *E_i_* (i = resource 1 or 2) and assimilation effort *ϕ* which determines the assimilation efficiencies for the two resources *A_i_* (Fig. 2). Resources are assimilated from the food proportionally to the available resource concentrations, the organism’s surface and the assimilation efficiency for either resource. Assimilates are then added to the reserves at rates *p_Ai_*. Reserves are consumed at rates *p_Ci_*, proportionally to their density. The synthesizing unit (SU) (Kooijman 1998) combines the two consumption fluxes at rate *p_SU_* which, after subtracting maintenance costs *p_M_*, results in structural growth of the organism. Structural volume is created with a constant density of both resources. If influxes of the SU are imbalanced excessive resources are rejected at rate *p_Ri_*, of which a fraction is recycled to the reserves while the rest is excreted. The resource density within the reserves is diluted by growth. Fluxes in and out of the reserves are expressed as resource density fluxes in units of resources per structural volume per time. We assume that the production of nutrient-specific digestive enzymes is energetically costly and that it involves trade-offs hindering the capacity of the consumer to maximize assimilation efficiency for all resources simultaneously (Zera & Harshman 2001). We therefore implemented a trade-off in the assimilation efficiency of the two co-limiting resources. Along this trade-off the organism allocates the assimilation effort (i.e. production of nutrient specific enzymes) towards the most limiting resource as a compensatory mechanism maintaining the nutrient uptake homeostasis (Clissold *et al*. 2010). Evidence suggests that in invertebrates such digestive processes are controlled by the nutrient status of the hemolymph (Bede *et al*. 2007). Accordingly, in the model, the allocation of assimilation effort is controlled by the balance of resources in the reserves. The adjustments of the assimilation effort are not immediate. We therefore implement a characteristic switching time that determines the speed at which an organism acclimates to changed resource conditions in the model (*see* Appendix 1, Fig. A1, A2 for further details). The model equations and parameter estimates are provided in Table 1. The resulting system of ODEs is numerically integrated with the *lsoda* solver from the *deSolve* package in *R* (R Development Core Team 2014) for the different treatments. Different parameter combinations, which represent limit cases of (i) no reserves, no variable assimilation, (ii) no variable assimilation, (iii) no reserves and (iv) the full model with reserves and variable assimilation are presented in Appendix 2, Fig. A3-4. We explore the model behavior in more detail and provide a sensitivity analysis for the three most influential parameters in Appendix 3 (Fig. A5-7).

**Table 1:** Model equations and parameter estimates. The model is loosely parametrized for *Daphnia*, when available, published values are given. All other values are set within reasonable biological ranges. ^a^ from (Vanoverbeke 2008); ^b^ inferred from (Koussoroplis *et al*. 2017b); ^c^ calculated from (Koussoroplis & Wacker 2016); ^d^ inferred from (Sperfeld & Wacker 2009) for cholesterol; ^e^ Authors’ personal observation on the *D. magna* clone used; ^f^ Slightly modified from AmP Daphnia magna version 2016/02/04 bio.vu.nl/thb/deb/deblab/add_my_pet/.

| State variables |  |  |  |  |
| --- | --- | --- | --- | --- |
| Symbol | Description | Differential equation | Initial condition | Unit |
| $\phi$ | Assimilation effort | $\frac{d\phi}{dt} = \frac{1}{\tau} (\phi_{Ti} - \phi)$ | 0.5 | 1 |
| $E_i$ | Reserve density | $\frac{dE_i}{dt} = p_{Ai} - p_{Ci} + \kappa p_{Ri} - (p_{SU} - p_M) E_i$ | $10^{-11}$ | mol mm<sup>-3</sup> |
| $V$ | Structural volume | $\frac{dV}{dt} = (p_{SU} - p_M) V$ | 0.008 | mm<sup>3</sup> |
| Functions |  |  |  |  |
| Symbol | Description | Equation |  |  |
| $\phi_T(x)$ | Target assimilation effort at reserve balance $x$ | $\phi_T(x) = \begin{cases} -\frac{1}{2}x^\gamma + 1, & x \leq 1 \\ \frac{1}{2}x^{-\gamma}, & x > 1 \end{cases}$ | | |
| $\varepsilon_i(\phi)$ | Assimilation efficiency | $\begin{aligned} \varepsilon_1 &= \phi^\alpha (\varepsilon_{max} - \varepsilon_{min}) + \varepsilon_{min} \\ \varepsilon_2 &= (1 - \phi)^\alpha (\varepsilon_{max} - \varepsilon_{min}) + \varepsilon_{min} \end{aligned}$ | | |
| Fluxes |  |  |  |  |
| Name |  | Equation | Unit |  |
| Assimilation flux | | $p_{Ai} = p_{Am} A_i n_i V^{-1/3}$ | mol mm<sup>-3</sup> d<sup>-1</sup> | |
| Catabolic flux | | $p_{Ci} = u E_i V^{-1/3}$ | mol mm<sup>-3</sup> d<sup>-1</sup> | |
| Outflow of $SU$ | | $p_{SU} = \frac{1}{\frac{1}{g_{max}} + \frac{1}{y_1 p_{C1}} + \frac{1}{y_2 p_{C2}} - \frac{1}{y_1 p_{C1} + y_2 p_{C2}}}$ | d<sup>-1</sup> | |
| Rejection flux | | $p_{Ri} = \begin{cases} p_{Ci} - p_{SU}/y_i, & p_{Ci} - p_{SU}/y_i > 0 \\ 0, & p_{Ci} - p_{SU}/y_i \leq 0 \end{cases}$ | mol mm<sup>-3</sup> d<sup>-1</sup> | |
| Maintenance flux | | $p_M = 0.3 \text{ d}^{-1} \text{ }^a$ | d<sup>-1</sup> | |
| Parameters |  |  |  |  |
| Symbol | Description | Value |  |  |
| $\tau$ | characteristic switching time of assimilation <sup>b</sup> | 0.1 d | | |
| $\alpha$ | curvature of assimilation trade-off | 3 | | |
| $\gamma$ | sensitivity for reserve balance | 5 | | |
| $n_i$ | resource concentrations in the food | $7 \times 10^{-3}$ mol/mol C (high cond.)<br>$3 \times 10^{-4}$ mol/mol C (low cond.) | | |
| $p_{Am}$ | maximum ingestion rate <sup>c</sup> | $4 \times 10^{-6}$ mol C mm<sup>-2</sup> d<sup>-1</sup> | | |
| $y_i$ | yield of structural volume per mole resource $i$ <sup>d</sup> | $1.5 \times 10^8$ mm<sup>3</sup>/mol | | |
| $g_{max}$ | maximum volume-specific growth rate <sup>e</sup> | 0.6 d<sup>-1</sup> | | |
| $\varepsilon_{max}$ | maximum assimilation efficiency | 0.9 | | |
| $\varepsilon_{min}$ | minimum assimilation efficiency | 0.1 | | |
| $u$ | Reserve conductance <sup>f</sup> | 1 mm d<sup>-1</sup> | | |
| $\kappa$ | fraction of resources rejected by the SU and returned to the reserves | 0.3 | | |

### Experiments

Stock cultures of *D. magna* were kept in ADaM medium (Kluttgen *et al*. 1994) at 20 °C with saturating amounts of the green algae *Acutodesmus obliquus* (SAG 276-3a, culture collection of algae, University of Göttingen, Göttingen, Germany). *Synechococcus elongatus* (SAG 89.79), a non-toxic cyanobacterium lacking essential sterols, was used as unialgal food in the experiments. P-replete *S. elongatus* (Syn P+) was cultured semi-continuously in aerated WC medium (Guillard & Lorenzen 1972) at dilution rates of 0.2 d^-1^. P-deficient *S. elongatus* (Syn P-) were cultivated without dilution using P-free WC medium. All cultures were kept at 20 °C at a light:dark cycle of 16:8 h. The details on determining C as well as P concentrations of Syn P- and Syn P+ cultures for the daily prepared food suspensions are published elsewhere (Lukas *et al*. 2011). By mixing Syn P+ and Syn P- cultures the appropriate P:C ratios of food suspensions were obtained. Cholesterol containing liposomes were produced following the protocol described in (Wacker & Martin-Creuzburg 2012) and supplemented to the food suspensions to obtain desired cholesterol concentrations.

Third-clutch juveniles of *D. magna* (< 24 h old) were born on *S. elongatus* of limiting resource conditions (1.25 mmol P molC^-1^, 0.25 μg cholesterol mgC^-1^). In the constant treatments daphnids experienced either constant high or low P (6.66 or 1.25 mmol molC^-1^) each in combination with constant high or low cholesterol (8; 0.25 µg mgC^-1^), or average conditions of high and low resource concentrations (3.95 mmol P molC^-1^; 4.125 µg cholesterol mg C^-1^). In the variance treatments high and low concentrations of either P or cholesterol fluctuated, while the other resource (cholesterol or P, respectively) was constant at high concentration. In the covariance treatments high concentrations coincided or alternated, respectively for positive and negative covariance. Three different fluctuation frequencies of the resource supply (1/24 h, 1/48 h and 1/96 h, for a total experimental duration of 96 h) were applied for variance and covariance treatments, where animals experienced either high or low concentrations of the respective varying resource(s) first (phase of fluctuation). Each treatment consisted of four replicates with six juvenile *D. magna* per replicate. The cholesterol variance experiment was performed twice and the data was pooled resulting in eight replicates. The experiments were conducted at 20°C in the dark, using glass beakers filled with 200 ml of food suspensions prepared with ADaM medium. Food suspensions of *S. elongatus* (2 mg C L^-1^) with targeted resource concentrations were prepared daily. Every 12 h (for 1/24 h fluctuation treatments) but at least every 24 h (for constant, 1/48 h and 1/96 h fluctuation treatments) daphnids were transferred into renewed food suspensions to avoid food quantity limitation and to simulate (co)variance in P and cholesterol. Initial dry mass (*M_0_*) of animals was determined at the beginning of the experiment using three subsamples of 10 juveniles dried for three days at 45°C and weighted on an electronic balance (± 1µg, CP2P; Sartorius, Göttingen, Germany). After four days the final dry mass (*M_t_*) of animals of each replicate was determined following the same procedure. Somatic growth rates g (d^-1^) were calculated as increase in dry mass using the equation

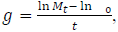

where *t* is the duration of the experiment in days. To test for effects of fluctuation frequency and phase (variance experiments) or fluctuation frequency and covariance direction (covariance experiment) on growth, for each experiment a two-way ANOVA was applied followed by Tukey HSD post-hoc tests. A nested-design two-way ANOVA was used to additionally test for effects of fluctuation phase nested within positive and negative covariance in the covariance experiment. Here, only the lowest fluctuation frequency of 1/96 h was considered. Means and associated 95% confidence intervals (CI) were estimated using a bootstrapping approach with 1000 repetitions (Efron & Tibshirani 1993). Regarding treatments of constant average resource supply 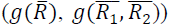 and non-linear averaging predictions for variance 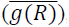, positive 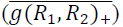, and negative covariance 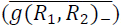, respectively, derived from constant resource condition treatments, a significant difference to observed growth rates under resource fluctuation was concluded in the absence of overlap between their respective CI.

## Results

### Model results

The simulated growth *g* of the consumer strongly changes with the fluctuation frequency of a resource *R* in the variance treatment, which confirms the temporal scale dependence. High resource fluctuation frequencies tend to be buffered by reserves and simulated growth converges towards the growth achieved under constant average resource conditions 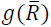 (Fig. 3A). However, as fluctuation frequency decreases, growth converges towards the non-linear average prediction 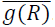. For a certain parameter range, growth under intermediate frequencies even becomes transiently lower than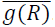 (Appendix 3, Fig. A8). The phase of fluctuation, i.e. starting with high or low resource concentration, also affects growth (Fig. 3A). The decrease of growth with decreasing fluctuation frequency is steeper when the organism experiences a low resource concentration first.

With covariance between the two resources *R_1_* and *R_2_* we also observe a strong dependence of consumer growth on the temporal scale of resource fluctuations (Fig. 4A). At low fluctuation frequencies the modelled growth trends towards 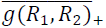 for positive and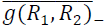 for negative covariance. As fluctuation frequency increases, simulated growth increases, whereas the covariance effect decreases. For positive covariance, the simulated growth saturates at 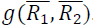. For negative covariance, we even observe growth rates higher than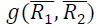, which results in an inversion of the covariance effect, i.e. fast fluctuations yield higher growth under negative rather the positive covariance. As we parametrized both resources identically, the phase of fluctuations does not affect growth in negative covariance scenarios. Only under positive covariance, an initially low resource supply decreases growth, which becomes most apparent at low fluctuation frequency (Fig. 4B).

In the model, we included three central concepts of nutritional physiology (Fig. 2). Dynamic acclimation allows the consumer to regulate the assimilation efficiencies for the two co-limiting resources according to the balance of stored resources compared to internal requirements along a trade-off curve (Appendix 1, Fig. A1-2). Reserves allow storage of the assimilated resources (Kooijman 2010). A synthesizing unit (SU (Kooijman 1998)) translates stored resources into structural growth. The results from the full model presented above may be better understood when the mechanisms are studied in isolation. The SU is indispensable and forms the null model as the two limiting resources always have to be combined to create new biomass. Only the inclusion of reserve, acclimation, or both into the model yields fluctuation frequency effects (Appendix 2, Fig. A3a-4a). Reserves increase growth across all fluctuation frequencies but most dominantly for high frequencies compared to the null model (Appendix 2, Fig. A3b-4b). They are thus responsible for the transition of the simulated growth from 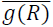 to 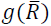 in the variance treatment and from 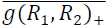 and 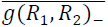 to 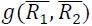 for positive and negative covariance, respectively. Also the effect of the fluctuation phase becomes pronounced when reserves are included (Appendix 2, Fig. A3b, Fig. A4b). Acclimation acts only when the two resources are fluctuating asynchronously, i.e. in the variance and negative covariance treatments (Appendix 2, Fig. A3c, Fig. A4c). For low fluctuation frequencies, acclimation and the resulting specialization on assimilating one resource increases growth compared to the null model. For high fluctuation frequencies, simulated growth trends towards the null model as acclimation becomes too slow and assimilation efficiencies remain in an intermediate, non-specialized range. Adding both, reserves and acclimation, to the null model increases growth rates even further at both low and high fluctuation frequencies and creates the inversion of the covariance effect (Appendix 2, Fig. A3d, Fig. A4d).

**Figure 3:**
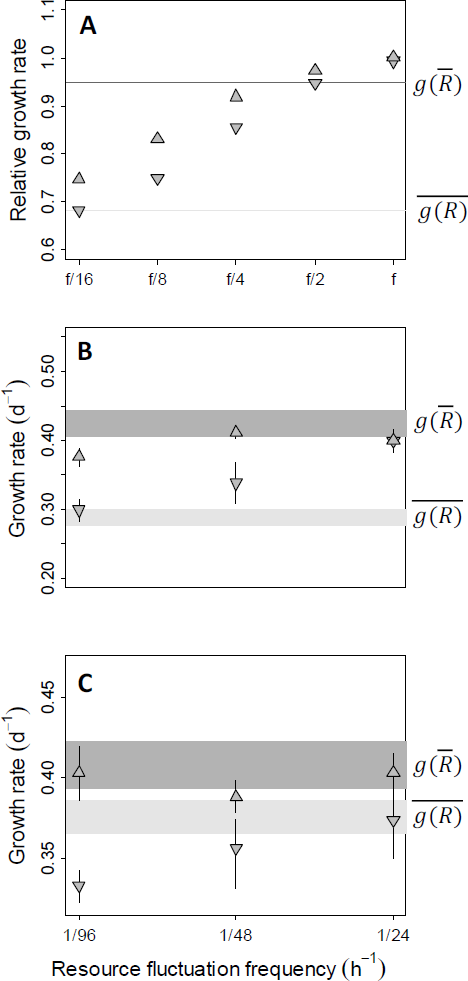
Effects of frequency and phase for single resource fluctuation. (**A**) Modelled relative somatic growth rates (fraction of the highest growth rate achieved) of a consumer exposed to fluctuating resource supply starting either with high (*upward triangle*) or low resource supply (*downward triangle*) at fluctuation frequencies relative to an arbitrary frequency *f*. The *darker horizontal line* is the growth under the same average constant resource supply, 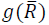, whereas the *lighter horizontal line* is the nonlinear averaging prediction of growth under variable resource supply, 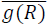 (see Fig.1a). (**B-C**)Observed juvenile growth rate of *Daphnia magna* (mean ± 95% C.I.) under (**B**) varying cholesterol or (**C**) phosphorus supply, while the other resource (phosphorus or cholesterol, respectively) is kept constant at saturating supply. The *darker shaded area* is the 95% confidence interval (C.I.) of 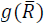The lighter shaded area is the 95% C.I. of 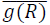.

### Experimental validation

We found a strong agreement between experimental results and the majority of predictions of the full model. In the experiment with fluctuating cholesterol supply, growth rates were affected both by fluctuation frequency (two-way ANOVA, F2,41 = 20.21, p < 0.001) and phase (F1,41 = 41.17, p < 0.001). As predicted, there were no growth differences between fluctuation phases at high frequencies (Fig. 3B), where growth rates were comparable to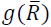 At low frequencies, however, growth rates were more reduced when exposed to low cholesterol concentrations first (interaction: F2,41 = 9.56, p < 0.001) and reached 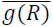 at the 1/96h frequency.

In the experiment with fluctuating P supply, growth rates were only marginally affected by fluctuation frequency (two-way ANOVA, F2,18 = 2.63, p = 0.10; Fig. 3C). In agreement with the model, growth rates were generally lower when animals experienced low P concentrations first (factor fluctuation phase, F1,18 = 33.86, p < 0.001). A marginal interaction between fluctuation frequency and phase (F2,18 = 3.01, p = 0.07) suggests the predicted increase in growth differences between the two phases at lower frequencies. When exposed to high P supply first, growth rates were high across all tested fluctuation frequencies and did not differ from 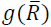(Fig. 3C). In contrast, when experiencing low P supply first, growth rates decreased with decreasing fluctuation frequency as predicted by the model, but were generally lower than 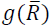 at all frequencies and even lower than 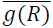 at the 1/96h frequency (Fig. 3C). Lower growth 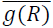 are also predicted by the full model for certain parameter ranges (Appendix 3, Fig. A8).

The experiments also confirmed the majority of model predictions for the two resource covariance scenarios (Fig. 4C). Similar to model predictions, frequency dependence was less pronounced for positive than for negative covariance, which is supported by a significant interaction of fluctuation frequency and covariance direction (F2,18 = 15.14, p < 0.001. As predicted, we observed an inversion of covariance effects; at the 1/96h frequency, negative covariance yielded a stronger growth decrease relative to 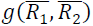 than positive covariance (Tukey HSD, p < 0.05), whereas at the 1/24h frequency the decrease was stronger for positive covariance (Tukey HSD, p < 0.05). The effects of fluctuation phase were tested for the 1/96 h frequency (Fig. 4D). Consistent with the model prediction, growth was affected by the fluctuation phase under positive covariance (nested ANOVA, F2,12 = 38.07, p < 0.001; Tukey HSD, p < 0.001), but was similar under negative covariance (Tukey HSD, p = 0.09).

**Figure 4:**
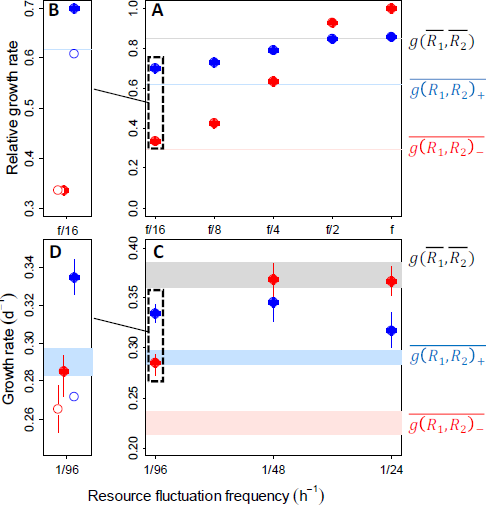
Effects of frequency, phase, and covariance for two co-limiting resources. (**A**) Modelled relative somatic growth rates (fraction of the highest growth rate achieved) of a consumer exposed to a positively (*blue*) or negatively (*red*) covarying supply in two co-limiting resources at fluctuation frequencies relative to an arbitrary frequency *f*. The *grey line* is the growth under the same average constant resource supply, 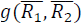, whereas the *red* and *blue* lines are the non-linear averaging prediction of growth under variable resource supply, 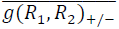 (see Fig.1B). (**B**) The effect of fluctuation phase: *Open blue circles*: starting with low resources conditions. (**C**) Observed juvenile growth rate of *Daphnia magna* (mean ± 95% C.I.) under positively or negatively covarying phosphorus and cholesterol supply. *Grey shaded area* is the 95% C.I. of 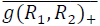. *Colored shaded areas* are the 95% C.I. of 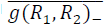 (red). (**D**) The effect of fluctuation phase at 1/96h.

## Discussion

Our study combines theoretical and experimental approaches to explore how variability in nutritionally complex environments influences the growth of a consumer limited by the supply of one or multiple limiting resources. For the first time, we explored the growth constraints imposed by covariance of two co-limiting resources. We found that the phase and the (co-)variance of fluctuating resource supply strongly constrain consumer growth and that this effect depends on the frequency at which the consumer experiences resource fluctuations.

Our modeling approach sheds light on the way resource acquisition and storage physiology can determine consumer performance in nutritionally fluctuating environments. The model incorporates three central physiological processes linked to nutrition: the ability to (i) optimize the acquisition of the most limiting resource, (ii) store resources, and (iii) combine resources at the required ratios into new biomass.

The adjustment of resource assimilation efficiency is a universal trait (Yin & Johnson 2000; Ashley *et al*. 2006; Karasov *et al*. 2011) enabling the organisms to maintain their biochemical homeostasis by focusing their acquisition efforts on the resource in short supply (Clissold *et al*. 2010; Wojewodzic *et al*. 2010; Karasov *et al*. 2011; Bansemer *et al*. 2016). In metazoans, this can be achieved by various mechanisms such as plasticity in gut transit, length, and mass, or in types and concentration of digestive enzymes (Karasov *et al*. 2011). Our acclimation model focuses on the case of digestive enzyme plasticity, which in *Daphnia* has been observed for gut proteases, lipases and phosphatases (Schwarzenberger *et al*. 2010; Wojewodzic *et al*. 2010; Koussoroplis *et al*. 2017b). This adjustment of digestive enzymes may involve two types of costs. First, modifying the pool of digestive enzymes requires energy, which is then not available for other processes such as growth or reproduction. Second, full acclimation requires time during which the consumer underperforms, since the shape of the assimilation efficiency trade-off prevents an optimal acquisition of the most limiting resource. Hence, the consumer’s growth may decrease below 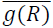. In the model simulations, when the resource fluctuations are slow enough (e.g. in a coarse grained environment), this delay becomes negligible compared to the time during which the organism benefits from full acclimation and specialization on the resource in short supply. As fluctuation frequency increases, the relative importance of the underperformance compared to the beneficial effect increases as the relative time spent at non-optimal assimilation efficiencies increases, thereby suppressing consumer growth. Eventually, the acclimation status will remain constant and adjust to the average nutritional conditions under which the organism assimilates both resources with suboptimal, reduced efficiency. In the model, acclimation is driven by imbalances of the resources in the reserves. Therefore, the underperformance, which indicates temporal costs for acclimation, occurs only when co-limiting resources are negatively covarying or under single resource variance.

Similar to acclimation, having reserves, i.e. a pool of assimilated resources stored in polymer form, is a universal trait in living organisms (Kooijman 2010). DEB theory assumes that assimilated resources first enter the reserves before being routed to growth and reproduction (Kooijman 2010). In our model reserves buffer high frequency fluctuations. This increases growth relative to the predictions for variable nutritional environments 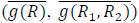 for generally observed performance curves (Fig. 1A-B). Thus, high frequencies enable the consumer to grow as well as in a constant nutritional environment of the same average quality 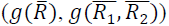, whereas under low frequencies performance converges towards 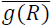 or 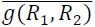 for variance and covariance scenarios, respectively (Koussoroplis *et al*. 2017a). The experimental observations are in agreement with the buffering effect. Including reserves in the model makes consumer growth depend not only on the frequency, but also on the phase of fluctuations, a pattern we also observed in our experiments. This indicates that in coarse-grained nutritional environments, where the temporal scale of resource fluctuations is large (i.e. low fluctuation frequency), timing is essential. Because initially accumulated reserves may be used during a subsequent phase of nutritional deprivation, experiencing a high quality nutritional environment creates a *window of opportunity* during early development and may dramatically improve performance (Malzahn & Boersma 2012).

We observed a strong covariance effect at low fluctuation frequencies. Growth under negative covariance is reduced compared to growth under positive covariance as the resource that is temporally in excess is rejected by the synthesizing unit and only partly recycled to the reserve compartment. Interestingly, combining reserves and acclimation dynamics (full model) cause an inversion in the direction of the covariance effect between high and low fluctuation frequencies, which also appears in our experimental results. Thus, not only the magnitude of the covariance effect but also its direction is determined by the resource fluctuation frequency. This illustrates the complexity of the challenges that consumers face when they need to acquire multiple resources in heterogeneous landscapes.

To tackle these challenges consumers may apply different physiological strategies, which become apparent in our model (see Appendix 2, Fig. A4b,c). An intriguing route to follow would be to study whether different types of environments might select for different strategies and if trade-offs between them exist. Reserve specialists, i.e. organisms with large reserves that are used up only slowly, could withstand periods of nutritional deprivation, and thus benefit in fast fluctuating environments. Assimilation specialists, i.e. organisms directing their assimilation efforts to specific, most limiting resources, would be well adapted to slowly fluctuating environments with sequential availability of limiting resources.

Our study confirms that nutritionally heterogeneous environments are a challenge for consumers because they involve various physiological costs (Hood & Sterner 2010; Wetzel *et al*. 2016; Wetzel & Thaler 2016; Wagner *et al*. 2017). Resource heterogeneity can decelerate invasion fronts (Giometto *et al*. 2017) or limit the success of agricultural pests (Underwood 2004) and is therefore suspected to regulate the dynamics of natural or agricultural ecosystems. Nutritional heterogeneity depends on various natural or anthropogenic factors. In agroecosystems for example, herbivore attacks on plants induce the production of volatile chemical signals that prompt chemical defenses in undamaged plants (Morrell & Kessler 2017). Such induced responses increase nutritional heterogeneity of individual plants or groups of aggregated plants for herbivores (Karban 2017). On the other side, anthropogenic landscape homogenization or reduced plant species diversity decreases the nutritional heterogeneity experienced by herbivores. However, any attempt to predict effects of such changes on herbivores is difficult as long as resource heterogeneity is ill-defined. In both examples above, the changes concern the resource variance affecting the fluctuation amplitude as well as the resource grain size in the landscape affecting the fluctuation frequency perceived by the consumer (Fig. 5a). Our results contribute major insights here, as they show that fluctuation amplitude and frequency may act on consumer growth in opposite directions: decreased heterogeneity due to lower amplitude increases consumer performance whereas decreased heterogeneity due to lower frequency decreases it (Fig. 5b-c). Hence, these two aspects of heterogeneity could compensate each other, e.g. a positive effect of landscape homogenization via the reduction of resource fluctuation amplitude could be weakened by a decrease in resource fluctuation frequency.

In nature, the physiological responses to environmental variability cannot be decoupled from behavior as these two aspects interact. Consumers permanently adjust their foraging behavior according to their nutritional status and their environment (e.g. resource peaks, predation risk, competitors or thermal preferences) (Kearney *et al*. 2010; Simpson & Raubenheimer 2012). Hence, even without a physical perturbation of the landscape, any alteration of the foraging behavior (e.g. staying longer in food patches, switching less often between complementary foods) (Lihoreau *et al*. 2017) modifies the perceived temporal structure of (co-)limiting resource supply. Understanding the linkages between foraging behavior, nutritional limitation and spatiotemporal distribution of (co-)limiting resources in the environment are clearly a fertile future field of research.

Given the multitude of factors shaping both the perceived variability of nutritional limitation as well as the physiological response of consumers, predicting their performance in a changing world is an intriguing yet challenging aspiration. Only an interdisciplinary approach allows to unravel the complex effects that may arise from the interplay of more frequent and drastic perturbations to the environment by extreme events on the one hand and anthropogenic environmental simplification on the other.

**Figure 5:**
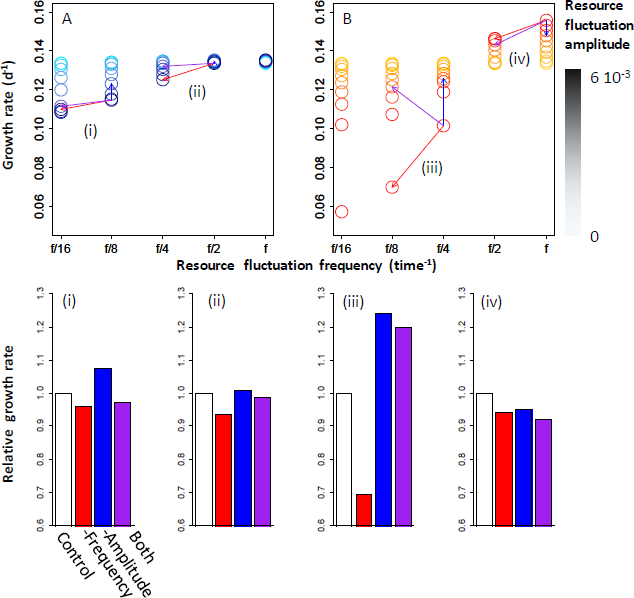
Combined effects of resource fluctuation amplitude and frequency. Modelled consumer somatic growth (**d**^-1^) for various combinations of resource fluctuation amplitude and frequency in the case of two positively (**A**) or negatively (**B**) covarying co-limiting resources. The colored arrows indicate various *scenarii* (i-iv) of decrease in either resource fluctuation amplitude (blue) or resource fluctuation frequency (red) alone or both (purple). The bar plots below illustrate the growth rate (relative to control) achieved in each of the scenarii.

## Acknowledgements

We thank Erik Sperfeld and Toni Klauschies for insightful comments on earlier versions of the manuscript. This study was supported by the German Research Foundation (KO5330/1-1 to AMK, WA2445/9-1 and WA2445/15-1 to AW)

## Appendix 1 – Assimilation trade-off

Organisms face the challenge of reconciling external resource availability and internal nutritional needs. This can be achieved by storing or egesting excess resources. An energetically more efficient route would be to prevent unnecessary ingestion in the first place and thus be able to spend that energy on the assimilation of resources in short supply. Here, we will present how we modelled the adaptive assimilation of resources that is driven by the internal resource requirements.

The internal requirements are set as the ratio of resource concentrations in the structural volume of the organism. The deviation between these requirements and the balance of resources in the reserves determines how much energy should be allocated to favour the assimilation of the more limiting resource over the other. The resource balance is given by 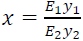 where *E_i_* are the reserve densities and *y*_*i*_ are the resource yields, i.e. the inverse resource concentrations. The resources are thus balanced for *x* = 1 and unbalanced otherwise.

Using an assimilation effort function *ϕ* this balance symmetrically translates into an estimate of whether energy investment should go into the assimilation of resource 1 (for *ϕ* → 1) or resource 2 (for *ϕ* → 0). The sensitivity for small imbalances *γ*, i.e. around *x* = 1, determines how drastic changes to the assimilation apparatus should be even for minor deviations (Fig. A1).

We assume that a trade-off exists between the assimilation efficiencies of the two resources, which can generally be exclusive, cooperative or inhibitory. An exclusive trade-off arises if energy or enzymes used to assimilate one resource lack to assimilate the other. If investments into assimilating one resource facilitate the assimilation of the other resource the trade-off is cooperative, e.g. if enzymes used for one resource are also used for the other. If for example cell surface limits the number of specific membrane receptors, an inhibitory trade-off would be expected.

The nature of the trade-off determines the shape of the trade-off curve, which in the model is set by the shape parameter *α* (Figure A2). If now the reserves become imbalanced the organism can change its target assimilation effort along this trade-off curve. The target assimilation effort is given by

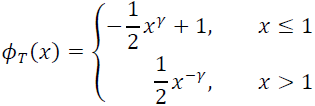

for reserve balance *x*. The assimilation effort then tracks the effort target in time.

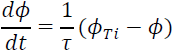

*τ* is the characteristic switching time of the assimilation apparatus and thus scales the speed at which the organism is able to acclimate.

**Figure A1:**
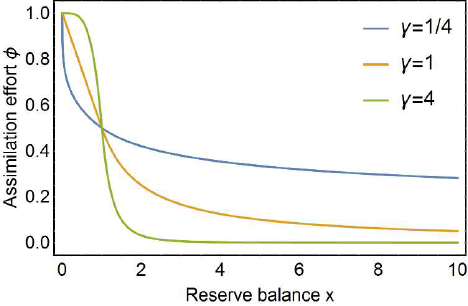
Assimilation effort. If reserves are highly unbalanced (*x* < 1 or *x* > 1) more assimilation effort is allocated to either one nutrient. If reserves are balanced (*x* = 1) both nutrients are assimilated equally. The sensitivity parameter *γ* gives the slope at *x* = 1. Throughout this study we assumed high sensitivity with *γ* = 4.

**Figure A2:**
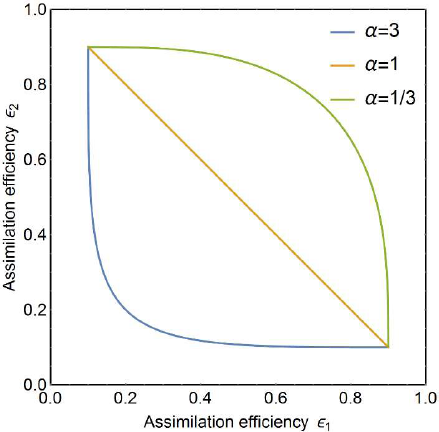
Shape of the assimilation trade-off curve. The trade-off curve determines the possible combinations of assimilation efficiencies. For *α* < 1 the assimilation trade-off is cooperative and at *α* = 1 it is exclusive. Throughout this study, we focused on inhibitory enzymatic assimilation pathways and thus used a shape parameter of *α* = 3.

## Appendix 2 – Detailed model results

To study the effect of reserves and acclimation in isolation and in concert, we created different model scenarios in which reserves and acclimation were switched on or off. In the simplest case (Null model, Figs. A3a and A4a) only the synthesizing unit (SU), which forms biomass from the two limiting resources, controls the model output. Reserves are excluded by increasing the reserve conductance 100-fold. Thus, the resources are not stored but directly passed on to the SU. Acclimation is excluded by fixing the assimilation efficiencies at the intermediate assimilation effort *ϕ* = 0.5 along the trade-off curve. The second scenario (Figs. A3b and A4b) combines SU and reserves by using the normal reserve conductance of *u* = 1 mm d^−1^. The output of the third model scenario (Figs. A3c and A4c) is shaped by SU and acclimation. The difference between the maximum and minimum assimilation efficiencies is increased and acclimation along the trade-off curve is possible. The parametrization of the full model scenario includes all three processes (Figs. A3d and A4d).

**Table A1:**
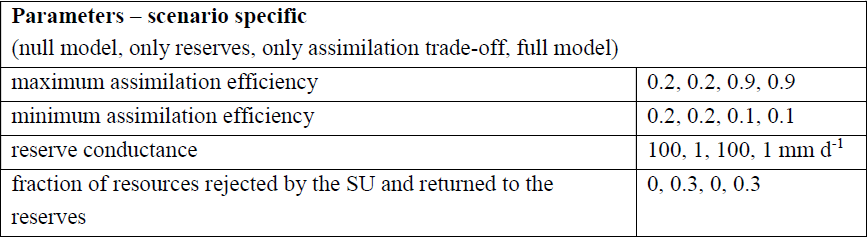
Parameters used in the different model scenarios.

| **Parameters – scenario specific**<br>(null model, only reserves, only assimilation trade-off, full model) |  |
| --- | --- |
| maximum assimilation efficiency | 0.2, 0.2, 0.9, 0.9 |
| minimum assimilation efficiency | 0.2, 0.2, 0.1, 0.1 |
| reserve conductance | 100, 1, 100, 1 $\text{mm d}^{-1}$ |
| fraction of resources rejected by the SU and returned to the reserves | 0, 0.3, 0, 0.3 |

**Figure A3:**
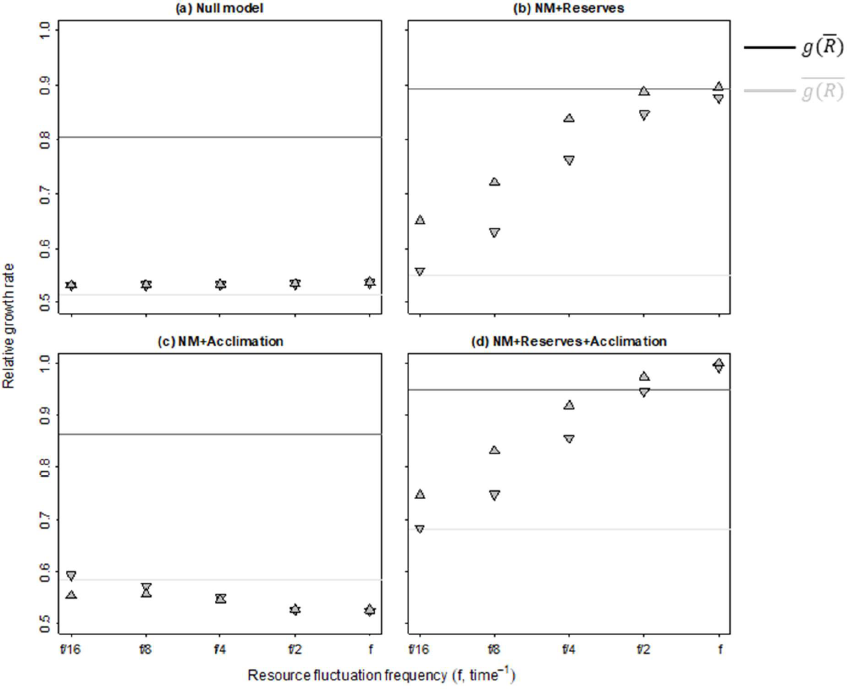
Results for the different model scenarios – One fluctuating nutrient. Predicted growth at constant resource concentration, 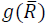 (dark grey lines), and non-linear averaging predictions for growth under variable resource concentration, 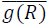 (light grey lines at fluctuation frequencies relative to an arbitrary frequency *f*. Simulation results for a single fluctuating resource and different phases, i.e. starting either with high (*upward triangle*) or low concentrations (*downward triangle*). (**a**) Only the synthesizing unit (SU) determines the growth rate. (**b**) Resources are stored in reserves before they are passed on to the SU. (**c**) The organism can acclimate its assimilation efforts depending on how balanced the nutrients in the reserves are compared to the nutrient requirements for growth. Nutrients however remain only very shortly in the reserves and are quickly passed on to the SU, i.e. no long-term storage is possible. (**d**) The organism is able to acclimate and also can store resources before they are passed on to the SU (Full model). Scenarios specific parameters are given in Tab. A1.

**Figure A4:**
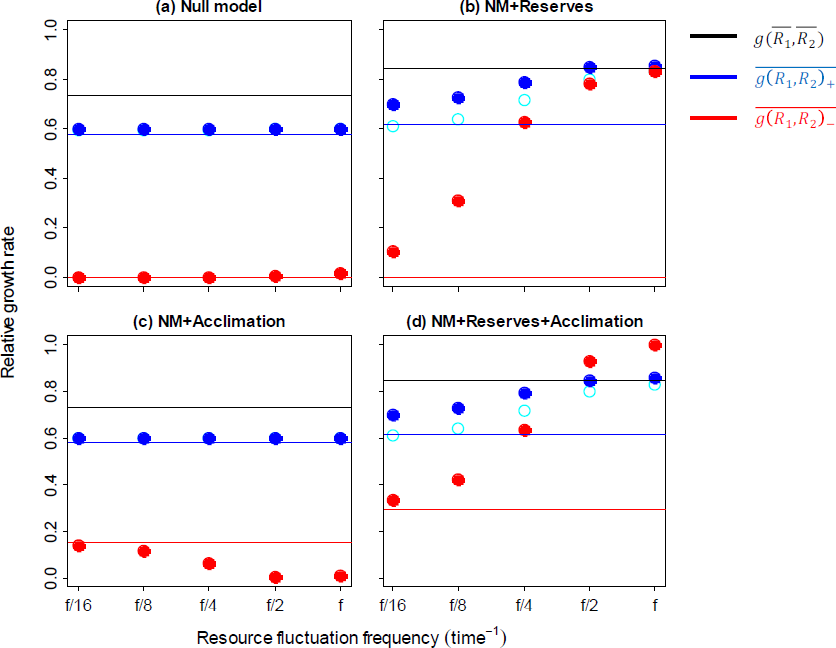
Results for the different model scenarios – Two fluctuating nutrients. Predicted growth at constant resource concentration, 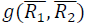 (black lines), and non-linear averaging predictions for growth under positively, 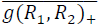 (blue lines) or negatively covarying resource concentrations, 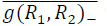 (red lines) at fluctuation frequencies relative to an arbitrary frequency f. Simulation results for positively (blue circles) or negatively (red circles) covarying resources and different phases, i.e. starting either with high (filled circles) or low concentrations (open circles). As the two nutrients are modelled identical, the phase has no effect under negative covariance. (**a**) Only the synthesizing unit (SU) determines the growth rate. (**b**) Nutrients are stored in reserves before they are passed on to the SU. (**c**) The organism can acclimate its assimilation efforts depending on how balanced the nutrients in the reserves are compared to nutrient requirements for growth. Nutrients remain only shortly in the reserves and are quickly passed on to the SU, i.e. no long-term storage is possible. (**d**) The organism is able to acclimate and also can store nutrients before they are passed on to the SU (Full model). Scenarios specific parameters are given in Tab. A1.

## Appendix 3– Sensitivity analysis

In this section, we will study the model behavior for the three most important parameters in the covariance treatments. These parameters for which our model shows the highest sensitivity are the characteristic switching time of assimilation *τ* (Fig. A5), the reserve conductance *u* (Fig. A6), and the shape parameter of the assimilation trade-off *α* (Fig. A7).

Overall, we see that the observed patterns are robust against changes in these parameters. The switching time *τ* sets the speed at which the assimilation effort can be changed. This acclimation is driven by the balance of reserves. *τ* does therefore only affect growth rates in variable resource conditions where the two resources do not co-vary positively (Fig. A5). Smaller *τ* allows for very fast changes of the assimilation effort allocation, which results in a sustained reserve balance. This increases growth under slowly fluctuating conditions as no resource becomes “too” limiting. For faster fluctuations, faster acclimation however decreases growth. Here, the resource fluctuations are already so fast that the reserves are not emptied within the phases of low supply of the respective resource and reserve imbalance is not hampering growth. Instead, for slower acclimation the assimilation efficiencies remain high for the nutrient that now also is supplied in high concentrations. This increases the overall nutrient assimilation and results in higher growth rates. This effect however decreases again at very fast resource fluctuations. The general patterns of increasing growth with increasing fluctuation frequency and growth rates above the predictions from constant conditions are present for a broad range of *τ*.

The reserve conductance *u* controls the outflow of resources out of the reserves towards the SU. A lower conductance thus decreases the overall resource flow to the SU and therefore also the growth rate under constant conditions, both for positive and negative covariance (Fig. A6). If resources are fluctuating, the reserves allow for storage of resources obtained during high resource phases, which then can be used during phases of low resources. Smaller conductances increase the storage capacity but also limit the supply rate to the SU. Thus, very small *u* may also reduce growth rates, but for slightly higher *u*, the resource provision to the SU is not the limiting factor. Here, a higher reserve capacity kicks in and increases growth for slow fluctuations (f/8 and f/4 in Fig. A6) compared to faster conductances, where this reserve capacity is smaller and the reserves are emptied during long periods of low external resources. If fluctuations are fast enough, reserve capacity is less important and growth rates are higher for higher conductances. Under positive covariance the growth rates at small conductance overshoot the predictions from constant conditions for low fluctuation frequencies if high concentrations are initially provided. We see here, that the effect of fluctuation phase becomes stronger for smaller *u* as here more of the initially obtained resources can be stored and used later. Again, the observed patterns are robust for broad parameter ranges in *u*. For fast fluctuations the growth rates trend towards the constant condition predictions under positive covariance and overshoot them for negative covariance.

The trade-off shape *α* defines the non-linearity of the assimilation trade-off. For *α* < 1 the second derivative of the trade-off curve is negative, zero for *α* = 1 and positive for *α* > 1. In Fig. A7 we see, that this non-linearity controls the inversion of the covariance effect. For *α* < 1 the trade-off curve is concave and the trade-off is weak. This results in on average higher assimilation efficiencies and increases growth, both under constant and variable conditions. Growth rates for positive and negative covariance converge for fast fluctuation frequencies. If *α* is large, the trade-off curve is convex and the trade-off is strong. This increases the effect of assimilation specialization towards one resource. On average, the assimilation efficiencies are smaller here, thus, growth rates are also generally lower. The specialization on the assimilation of one resource increases the total resource assimilation if the resources are provided sequentially. Thus, we observe here the inversion of the covariance effect and growth rates become higher for negative than for positive covariance at fast fluctuation frequencies.

For certain combinations of *τ* and *u* we observe a modelled growth rate below the predictions from non-linear averaging (Fig. A8), especially for large *τ*, i.e. slow acclimation of the assimilation efficiencies, and large *u*, i.e. small storage capacity. Here, a mixture of mal-adaptation and lacking compensation by storage causes an underperformance of the organism under fluctuating conditions relative to the predictions from non-linear averaging.

**Figure A5:**
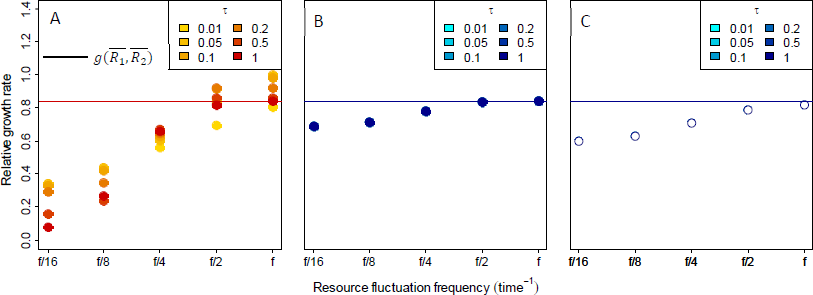
Full model sensitivity to the characteristic switching time τ. (**A**) Negative covariance. (**B**) Positive covariance and starting with high resources. (**C**) Positive covariance and starting with low resources. All growth rates are rescaled by their maximum.

**Figure A6:**
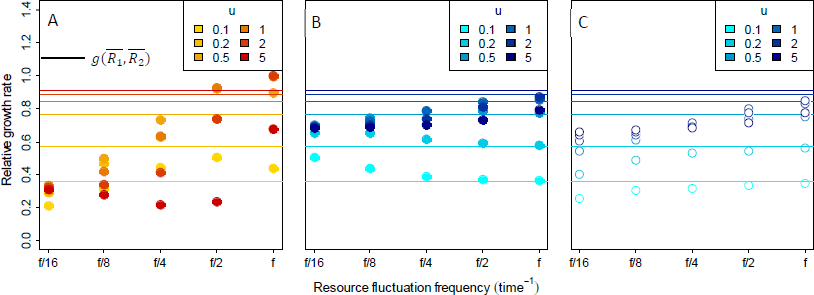
Full model sensitivity to the reserve conductance *u*. (**A**) Negative covariance. (**B**) Positive covariance and starting with high resources. (**C**) Positive covariance and starting with low resources. All growth rates are rescaled by their maximum.

**Figure A7:**
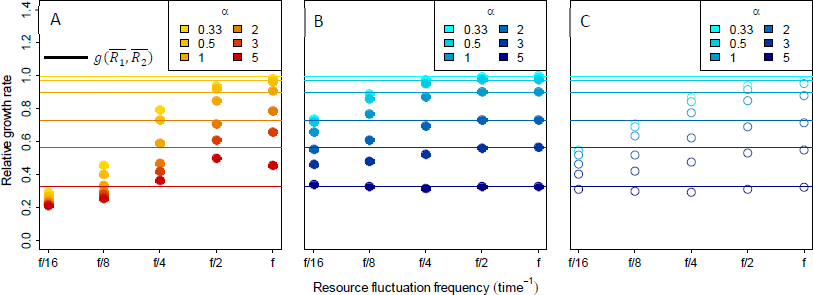
Full model sensitivity to the trade-off shape α. (**A**) Negative covariance. (**B**) Positive covariance and starting with high resources. (**C**) Positive covariance and starting with low resources. All growth rates are rescaled by their maximum.

**Figure A8:**
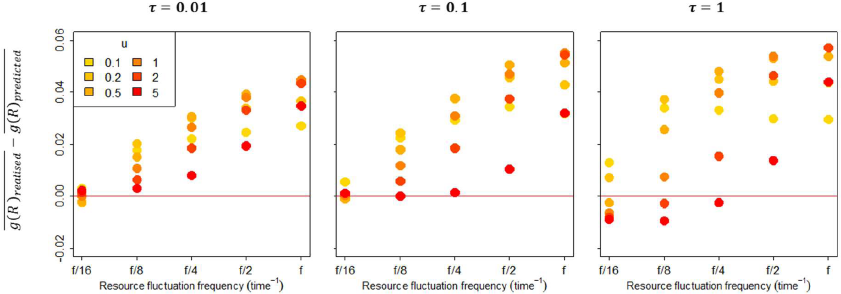
Transient underperformance in the full model: sensitivity to reserve conductance *u* and characteristic switching time τ. Single resource fluctuation starting with low resources. Underperformance is defined as the situation where modelled growth rate under fluctuating conditions, 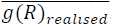is lower than the growth predicted by non-linear averaging, 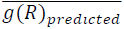.

